# The Impact of Restriction-Modification Systems on Mating in *Haloferax volcanii*

**DOI:** 10.1101/2020.06.06.138198

**Authors:** Matthew Ouellette, Andrea M. Makkay, Artemis S. Louyakis, Uri Gophna, J. Peter Gogarten, R. Thane Papke

## Abstract

Halobacteria have been observed to be highly recombinogenic, frequently exchanging genetic material. Several barriers to mating in the Halobacteria have been examined, such as CRISPR-Cas, glycosylation, and archaeosortases, but these are low barriers that do not drastically reduce the recombination frequency. Another potential barrier could be restriction-modification (RM) systems, which cleave DNA that is not properly methylated, thus limiting the exchange of genetic material between cells which do not have compatible RM systems. In order to examine the role of RM systems on limiting recombination in the Halobacteria, the impact of RM systems on cell-to-cell mating in *Haloferax volcanii*, a well-characterized method of genetic exchange and recombination in a halobacterial species, was examined. Strains which possessed all naturally-occurring RM system genes in *H. volcanii* (RM^+^) and strains without these RM systems (ΔRM) were mated together to compare the efficiency of gene transfer between RM-compatible strains and RM-incompatible strains. The results indicated that mating RM-incompatible strains together resulted in a decrease in gene transfer efficiency compared to mating RM-compatible strains together, suggesting that RM systems limit mating in *H. volcanii*, but do not act as absolute barriers to recombination. Therefore, RM systems are low barriers to recombination in the Halobacteria, with RM-incompatible strains exchanging genetic material at a lower frequency than those with compatible RM systems, similar to other low recombination barriers in the Halobacteria.

## 1. Introduction

In Bacteria and Archaea, distantly related organisms can exchange genetic material through horizontal gene transfer (HGT) and recombination. Many different strategies exist to allow for horizontal transfer of genetic material (Blakely, 2015). One method is natural transformation, where extracellular DNA is acquired via natural competence systems (Chen and Dubnau, 2004). Gene transfer can also occur via transduction, where bacteriophages transfer DNA between host cells (Touchon *et al*., 2017). Cells can also transfer genetic material via conjugation, where cells come into contact with each other and transfer plasmid DNA and integrative conjugal elements between each other, usually through specific structures such as a type IV pilus (Banuelos-Vazquez *et al*., 2017). Newly acquired genetic material which is not self-replicating is incorporated into the host’s genome via homologous recombination, in which the DNA is integrated at homologous sites in the genome (Rocha *et al*., 2005). Although gene transfer can occur between species of distant lineages, barriers to exchange do exist which can prevent transfer between certain species. Some of these barriers are physiological in nature (Thomas and Nielsen, 2005), such as surface exclusion which limits conjugation by preventing pilus formation and DNA transfer between the cells (Arutyunov and Frost, 2013), host range limitations of transferred plasmids (Hulter *et al*., 2017), or the lack of DNA uptake signals in eDNA which prevents its use by naturally competent cells (Smith *et al*., 1999; Spencer-Smith *et al*., 2016). Barriers to recombination can result in HGT being more likely to occur among more closely related strains and species over transfer events between more distantly-related species (Andam and Gogarten, 2011).

Restriction-modification (RM) systems can also potentially act as barriers to gene transfer. RM systems consist of a restriction endonuclease (REase) and a DNA methyltransferase (MTase) which both recognize the same target sequence of DNA. The MTase will methylate a base at the target site, whereas the REase will cleave the site if it is not methylated (Ershova *et al*., 2015). These systems act as defense mechanisms for their host organisms, in which potentially harmful foreign DNA which is not properly methylated is cleaved by the REase while the host’s own genome is protected due to methylation (Bickle and Kruger, 1993; Tock and Dryden, 2005). The ability of these systems to restrict foreign DNA could allow them to limit genetic exchange between species, thus potentially driving the diversification of microbial populations (Erwin *et al*., 2008; Budroni *et al*., 2011). Studies have demonstrated that RM systems can limit conjugal transfer of plasmids (Roer *et al*., 2015), and that they can limit the size of recombinant DNA fragments that are obtained via natural transformation (Lin *et al*., 2009).

Genetic recombination has been observed to occur frequently in several representatives of the halophilic archaeal class Halobacteria. In a study by Papke *et al*. (2007), *Halorubrum* strains isolated from saltern ponds in Spain and a hypersaline lake in Algeria were observed to cluster (<1% DNA sequence divergence for five housekeeping genes) into three major phylogroups, with sequence diversification being driven primarily by recombination within the phylogroups rather than mutations. In a study by Fullmer *et al*. (2014), *Halorubrum* isolates from a hypersaline lake in Iran were observed to cluster into distinct phylogroups, with each group sharing an average nucleotide identity (ANI) of greater than 98%. Recombination was also observed to occur frequently within the phylogroups, but at a lower rate between the phylogroups (Fullmer *et al*., 2014). Because recombination is more frequent between closely-related *Halorubrum* spp. strains, and less so with more distantly related species, barriers to gene flow limit genetic exchange, possibly resulting in genetically isolated populations and the diversification of haloarchaeal populations.

One mechanism of gene transfer in the Halobacteria is cell-to-cell mating, which has been characterized in *Haloferax volcanii* (Rosenshine *et al*., 1989). In this process, the cells come together and fuse into a heterodiploid state which contains the genetic material of both parental cells. This state allows for gene transfer and recombination between the parental cells. After genetic exchange, the cells will separate into hybrids of the parental strains (Rosenshine *et al*., 1989; Ortenberg *et al*., 1998; Naor *et al*., 2012). This process appears to have a low species barrier. In a study by Naor *et al*. (2012), *H. volcanii* was mated with the closely-related species *Haloferax mediterranei*, resulting in successful hybrids. However, the interspecies mating efficiency was observed to be lower than intraspecies mating events, suggesting that barriers to recombination exist which limit interspecies mating events between *H. volcanii* and *H. mediterranei*. One barrier to mating is CRISPR-Cas systems, which consist of short, repeated, spacer sequences acquired from foreign genetic elements that act as immunity systems for the host (Barrangou *et al*., 2007). These spacer sequences are used to produce RNAs known as crRNAs, which interact with Cas proteins to target and degrade invasive, foreign elements (Koonin *et al*., 2017). CRISPR-Cas systems have been identified in halobacterial species such as *H. volcanii* and *H. mediterranei* (Li *et al*., 2013; Maier *et al*., 2019), and research has indicated that these systems can limit interspecies mating between *H. volcanii* and *H. mediterranei* when the chromosome of one species is designed to be targeted by the partner species, although they do not act as total barriers to recombination (Turgeman-Grott *et al*., 2019). The glycosylation of surface glycoproteins has also been observed to affect mating in *H. volcanii*. A study by Shalev *et al*. (2017) tested the mating efficiency of *H. volcanii* mutant with the glycosylation genes *aglB* and *agl15*, and observed a dramatic decrease in mating efficiency when the deletion mutants were mated together, indicating that proper glycosylation is required for mating. However, mating the deletion mutant with their parental strains resulted in a less notable decrease in mating, suggesting that glycosylation limits mating between strains with different glycosylation patterns, but is likely not an absolute barrier to recombination (Shalev *et al*., 2017). Modification of surface proteins by archaeosortases has also been observed to affect mating. A study by Abdul Halim *et al*. (2013) examined the mating efficiency of a *H. volcanii* strain with the archaeosortase gene *artA* deleted, and observed that mating decreased in the deletion mutant compared to the parental strain, but was not a total barrier to recombination. Overall, CRISPR-Cas, glycosylation, and archaeosortases act as low barriers to mating, but do not prohibit recombination completely.

Another possible barrier to mating in the *H. volcanii* might be RM systems. Studies have characterized a few of these systems in *H. volcanii*, including a Type I RM system which targets the motif GCABN_6_VTGC (Ouellette *et al*., 2015; Ouellette *et al*., 2018). A Type IV REase known as Mrr has also been characterized in *H. volcanii*, and has been observed to reduce transformation efficiency on GATC-methylated plasmids (Holmes *et al*., 1991; Allers *et al*., 2010). However, the overall role of these systems on cell-to-cell mating and recombination has not been examined in detail. In this study, derivatives of the *H. volcanii* RM system deletion strain from Ouellette *et al*. (2018) were used in mating experiments to determine the impact of RM systems on cell-to-cell mating in *H. volcanii*.

## 2. Materials and Methods

### 2.1. Strains and Growth Conditions

All strains and plasmids used in this study are recorded in Table 1. *Haloferax volcanii* strains were grown in either rich undefined medium (Hv-YPC) or selective undefined medium (Hv-Ca) developed by Allers *et al*. (2004) and listed in the *Halohandbook* (Dyall-Smith, 2009). Media was supplemented with uracil (50 µg/mL) and 5-fluoroorotic acid (5-FOA) (50 µg/mL) as needed to grow *ΔpyrE2* strains. For *ΔtrpA* strains, the media was supplemented with tryptophan (50 µg/mL) as needed, whereas thymidine (40 µg/mL) and hypoxanthine (40 µg/mL) were supplemented as needed when growing *ΔhdrB* strains. The strains were grown at 42 °C while shaking at 200 rpm. *Escherichia coli* strains were grown at 37 °C while shaking at 200 rpm in lysogeny broth (LB), with ampicillin (100 µg/mL) added to the medium as needed.

**Table 1.** List of plasmids and strains used in this study.

| Strain/Plasmid Name | Description | Source |
| --- | --- | --- |
| *H. volcanii* H53 | ΔpyrE2 ΔtrpA uracil and tryptophan auxotrophic strain of wild-type *H. volcanii* | (Allers *et al.*, 2004) |
| *H. volcanii* H98 | ΔpyrE2 ΔhdrB uracil, thymidine, and hypoxanthine auxotrophic strain of wild-type *H. volcanii* | (Allers *et al.*, 2004) |
| *H. volcanii* ΔRM | Deletion strain of RM genes *HVO_0794*, *rmeRMS*, *HVO_A0006*, *HVO_A0074*, *HVO_A0079*, and *HVO_A0237* derived from a ΔpyrE2 uracil auxotrophic strain of *H. volcanii* | (Ouellette *et al.*, 2018) |
| *H. volcanii* RM ΔtrpA | ΔtrpA tryptophan auxotrophic strain derived from ΔRM | This study |
| *H. volcanii* RM ΔhdrB | ΔhdrB thymidine and hypoxanthine auxotrophic strain derived from ΔRM | This study |
| *E. coli* HST08 | *E. coli* cloning strain | Clontech, Cat. # 636763 |
| pTA95 | Plasmid used to delete *trpA* gene | (Allers *et al.*, 2004) |
| pTA155 | Plasmid used to delete *hdrB* gene | (Allers *et al.*, 2004) |

### 2.2. *Deletion of* trpA *and* hdrB *Genes from ΔRM Strain*

Plasmids pTA95 and pTA155 were used to delete *trpA* and *hdrB*, respectively, from *H. volcanii* strain ΔRM. These plasmids were transformed into the ΔRM strain using the polyethylene glycol (PEG)-mediated transformation protocol from the *Halohandbook* (Dyall-Smith, 2009), with resulting transformants being plated on Hv-Ca for 5-7 days. Screening for pop-ins was performed via colony PCR with screening primers (Table 2) and gel electrophoresis for visualization. Pop-outs were obtained by plating confirmed pop-ins on Hv-Ca plates with 5-FOA, uracil, tryptophan, thymidine, and hypoxanthine. Successful pop-outs were identified via replica plating onto Hv-Ca plates with uracil but without tryptophan, thymidine, or hypoxanthine, as well as via the colony PCR screening method used to detected pop-ins.

**Table 2.**
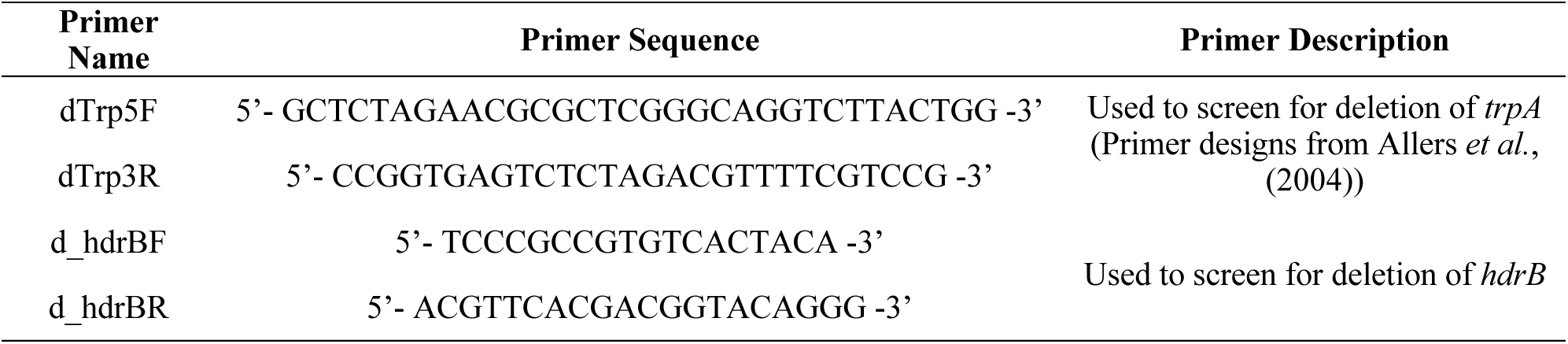
List of primers used in this study.

### 2.3. *Mating H53 and H98 Strains with RM* ΔtrpA *and RM* ΔhdrB *Strains*

Experiments were set up following a mating protocol adapted from Naor *et al*. (2012). Cultures of H53, H98, RM *ΔtrpA*, and RM *ΔhdrB* were grown in triplicate across three experiments (9 replicates total) to an OD_600_ of ∼1-1.1, with 2 mL of two different cultures applied to 0.2µm filters. The cultures were mixed in the following combinations: H53 × H98, H53 × RM *ΔhdrB*, RM *ΔtrpA* × H98, and RM *ΔtrpA* × RM *ΔhdrB*. The resulting filters were then placed on plates of Hv-Ca with uracil, tryptophan, thymidine, and hypoxanthine and incubated at 42 °C for 2 days. The filters were then transferred to 2-mL tubes containing 1 mL of liquid Hv-Ca medium and shaken at 200 rpm for ∼1 hour to resuspend the cells in the medium. The resuspended cultures were then diluted and plated onto Hv-Ca plates with uracil to determine the number of recombinants, and Hv-Ca plates with uracil, tryptophan, thymidine, and hypoxanthine to determine the total number of viable cells. The plates were incubated at 37 °C for 1 week. The number of recombinants was divided by the total number of viable cells to calculate the mating efficiency of each mated combination. Mann-Whitney U tests were performed in R with the package ggpubr v0.2.1 (Kassambara, 2018). A boxplot was constructed in R using the package ggplot2 v3.2.0 (Wickham, 2016).

## 3. Results

### Mating Efficiency is Lower When Mating RM-incompatible Strains

In order to determine whether RM systems in *H. volcanii* can act as a barrier to mating, *H. volcanii* strains H53 and H98, which contained the full set of RM system genes (RM^+^), and ΔRM derivative strains which were missing RM system genes (RM *ΔtrpA*, RM *ΔhdrB*) were mated together (Figure 1). Each strain was mated with a partner strain with a matching set of RM genes (RM-compatible; H53 × H98, RM *ΔtrpA* × RM *ΔhdrB*) and a partner strain without matching RM gene sets (RM-incompatible; H53 × RM *ΔhdrB*, RM *ΔtrpA* × H98). In each mating event, one strain is a tryptophan auxotroph (*ΔtrpA*) and the other strain is a thymidine auxotroph (*ΔhdrB*). Therefore, by mating the strains together and plating them on selective plates without tryptophan or thymidine, recombinants can be selected which contain both *trpA* and *hdrB* from both parental strains, and mating efficiency can be determined by calculating the number of colonies on the selective plates divided by the total number of viable cells for each mating event.

**Figure 1.**
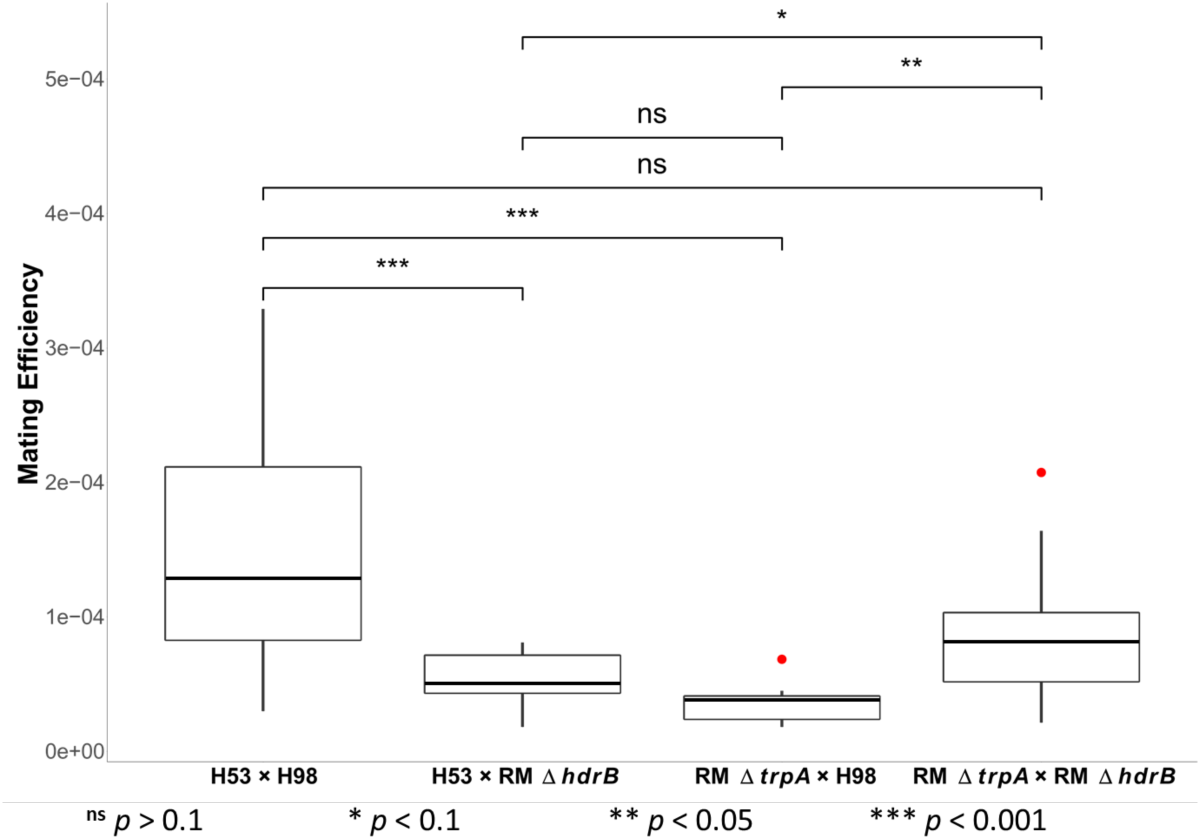
Box plot for mating efficiencies of *H. volcanii* strains H53 and H98 with RM system genes (RM+) and ΔRM derivative strains without RM systems (RM *ΔtrpA*, RM *ΔhdrB*). Mating crosses between RM-compatible strains (H53 × H98, RM *ΔtrpA* × RM *ΔhdrB*) and RM-incompatible strains (H53 × RM *ΔhdrB*, RM *ΔtrpA* × H98) were performed. Mating efficiency is expressed as the average number of colonies on selective plates divided by the total number of viable cells of each mating cross performed in triplicate for three experiments (9 replicates total). Red dots represent outliers. The *p*-values are from Mann-Whitney U tests of the differences between each mating cross.

The results indicate that mating H53 with H98 (H53 × H98) had an average mating efficiency of 1.6 × 10^−4^. In comparison, mating H53 with RM *ΔhdrB* (H53 × RM *ΔhdrB*) resulted in an average mating efficiency of 5.2 × 10^−5^, representing a ∼66% decrease from H53 × H98. The difference between these efficiencies was supported as significant by Mann-Whitney U (*p* = 0.004). When mating RM *ΔtrpA* with H98 (RM *ΔtrpA* × H98), the average mating efficiency was 3.7 × 10^−5^, representing a ∼77% decrease from H53 × H98. This was a significant difference supported by Mann-Whitney U (*p* = 0.001).

When mating RM *ΔtrpA* with RM *ΔhdrB* (RM *ΔtrpA* × RM *ΔhdrB*), the average mating efficiency was observed to be 9.2 × 10^−5^. This was a ∼43% decrease in mating efficiency from H53 × H98, but this difference was not significant according to Mann-Whitney U (*p* = 0.14). The mating efficiency increased from H53 × RM *ΔhdrB* by ∼70%. However, the difference was not strongly supported as significant by Mann-Whitney U (*p* = 0.09). The mating efficiency of RM *ΔtrpA* × RM *ΔhdrB* increased by ∼149% from RM *ΔtrpA* × H98, and the differences were supported as significant via Mann-Whitney U (*p* = 0.02). A difference was also observed between the mating efficiency of the RM-incompatible mating events, with RM *ΔtrpA* × H98 exhibiting a ∼33% lower mating efficiency than H53 × RM *ΔhdrB*. However, this difference was not supported as significant by Mann-Whitney U (*p* = 0.11). Overall, the results indicate that mating efficiency between RM-compatible strains is higher than the mating efficiency between RM-incompatible strains in *H. volcanii*.

## 4. Discussion

This study provides evidence that RM systems might act as post-fusion barriers to recombination in *H. volcanii*. When RM-compatible strains were mated together, such as H53 with H98 and RM *ΔtrpA* with RM *ΔhdrB*, the recombination efficiencies were similar to each other. The average mating efficiencies when crossing the RM-compatible strains were both close to the 1 × 10^−4^ intraspecies mating efficiency for *H. volcanii* observed by Naor *et al*. (2012), with H53 × H98 having an average mating efficiency of ∼1.6 × 10^−4^ and RM *ΔtrpA* × RM *ΔhdrB* having an average mating efficiency of ∼9.2 × 10^−5^. However, when RM-incompatible strains were mated together, such as H53 × RM *ΔhdrB* or RM *ΔtrpA* with H98, the mating efficiencies were observed to be lower than mating between RM-compatible strains. This difference indicates that when cells do not have compatible sets of RM systems, they recombine less frequently, likely due to the RM systems from RM^+^ cells cleaving DNA from ΔRM partner cells which is not properly methylated at recognition sites of the RM systems, thus preventing recombination from occurring.

RM systems have been observed to limit conjugation events in bacteria. In a study by Roer *et al*. (2015), for example, conjugation was observed to be limited in *E. coli* by the RM system EcoKI when using plasmids with unmethylated EcoKI target sites, although they are not a major barrier to conjugal transfer. A ∼85% reduction in transfer rate was observed when unmethylated plasmids were transferred into a recipient strain with an active EcoKI system in comparison to the rate when using a recipient strain with a deactivated EcoKI system (Roer *et al*., 2015). This reduction in *E. coli* conjugation is slightly higher than the ∼66-77% decrease in mating efficiency observed when mating RM-incompatible strains together, suggesting that RM systems have a slightly lower impact on cell-to-cell mating in *H. volcanii* than they do on conjugation in *E. coli*.

A few studies have also suggested that RM systems can limit recombination as well. Phylogenetic studies in *Haemophilus influenzae* and *Neisseria meningitidis* have suggested RM systems could act as barriers to recombination and drive population diversification (Erwin *et al*., 2008; Budroni *et al*., 2011). A Type III RM system in *Staphylococcus aureus* has also been demonstrated to prevent natural transformation on DNA from other bacterial species such as *E. coli* (Corvaglia *et al*., 2010). However, other studies have indicated that DNA cleaved by RM systems are still able to recombine, and that RM systems only limit the size of recombined DNA fragments (Vasu and Nagaraja, 2013). A study by Chang and Cohen (1977) indicated that the REase of RM system EcoRI was able to mediate site-specific recombination in *E. coli*. A study of natural transformation in *Helicobacter pylori* by Lin *et al*. (2009) indicated that RM systems limited the size of fragments imported during transformation, but did not prevent recombination of those fragments. The decreased mating between RM-incompatible strains observed in this study suggests that RM systems act to reduce recombination in *H. volcanii* rather than just limit fragment size of DNA during mating, and might be more effective if the methylation site were highly distributed around the chromosome or in the middle of our genetic markers.

The results also indicate that the RM *ΔtrpA* and RM *ΔhdrB* strains, when mated together, had a slightly decreased mating efficiency compared to when H53 and H98 were mated together. This result suggests that strains derived from ΔRM, which are missing active RM systems, are less efficient at mating than RM^+^ strains H53 and H98, which would indicate the RM systems have an important role in facilitating recombination. However, this difference was not strongly supported by Mann-Whitney U, so it is possible that this difference was due to chance. Also, the observation of an increase in mating efficiency when compared to H53 × RM *ΔhdrB* and RM *ΔtrpA* × H98 suggests that, even if the ΔRM strains themselves have a lower mating efficiency, RM-incompatibility between strains also results in reduced mating efficiency.

In *H. volcanii*, there are two major methylated motifs throughout the genome: the m4C motif C^m4^TAG and the m6A motif GCAm6BN_6_VTGC (Ouellette *et al*., 2015). The Cm4TAG motif is methylated by the orphan MTase HVO_0794 and the GCAm6BN_6_VTGC motif is methylated by the Type I RM system RmeRMS (Ouellette *et al*., 2018). Since HVO_0794 is an orphan MTase and is not associated with a RM system, it is unlikely to have an impact on mating efficiency and recombination. The most likely candidate for reducing mating between RM-incompatible strains in *H. volcanii* is the RmeRMS system. In the ΔRM strain and its derivatives, the genome is missing the operon that encodes the RmeRMS system, and is unmethylated at its target sites (Ouellette *et al*., 2018). Therefore, the target sites of RmeRMS would be exposed to cleavage when ΔRM derivative strains are mated with strains H53 or H98, which have the RmeRMS system. While *H. volcanii* also has the Type IV REase Mrr, which limits transformation efficiency with GATC-methylated plasmids from *E. coli* (Holmes *et al*., 1991; Allers *et al*., 2010), this REase has not been demonstrated to cleave methylated *H. volcanii* DNA and is unlikely to affect mating and recombination. Therefore, the decrease in mating between RM-incompatible strains is likely the result of RmeRMS in the RM^+^ strain cleaving unmethylated sites from the ΔRM strain and, therefore, limiting recombination between the strains. However, examination of the *trpA* and *hdrB* marker gene sequences, which are exchanged during mating, indicated that there are no RmeRMS sites located within those genes. The closest RmeRMS site to *trpA* is located 897 bp upstream of the gene, and the closest site to *hdrB* is located 6441 bp upstream of the gene.

This would suggest that RmeRMS could negatively impact recombination even when the restriction sites are distantly located from the genes of interest. One possible explanation is that RM-incompatibility between strains reduces the time that the cells can coexist after fusing together due to DNA cleavage, resulting the cells separating early from each other and, therefore, reducing chances for recombination between the cells. It is possible that, if the restriction sites were located closer to the genes of interest, or within the genes themselves, the mating efficiency between the RM-incompatible strains would decrease further. Since the RmeRMS site is located closer to *trpA* than *hdrB*, it is possible that recombination is more limited for *trpA* than *hdrB*. This may explain the difference in mating efficiency observed between the two RM-incompatible mating crosses. However, this difference was not supported as statistically significant. Future mating experiments using ΔRM derivative strains complemented with the *rmeRMS* operon could confirm the role of this RM system in limiting mating in *H. volcanii*.

RM-incompatibility may also be an explanation for the lower interspecies mating efficiency observed between *H. volcanii* and *H. mediterranei* (Naor *et al*., 2012). According to the RM database REBASE (Roberts *et al*., 2015), *H. mediterranei* has only three RM-related genes (http://rebase.neb.com/cgi-bin/onumget?8920): a Type IV REase (Mrr), an orphan m4C CTAG MTase (M.Hme33500I), and an orphan m4C MTase which modifies the motif HGC^m4^WGCK (M.Hme33500II), also described recently (DasSarma *et al*., 2019). Since it has no Type I RM system analogous to RmeRMS in *H. volcanii*, its genome is unmethylated at the target sites of that RM system. Therefore, the genome of *H. mediterranei* is exposed to cleavage by the RmeRMS system when mating with *H. volcanii*, which could limit recombination. Interestingly, the interspecies mating efficiency observed by Naor *et al*. (2012) was 4.2 × 10^−5^, which is between the mating efficiencies observed when crossing RM-incompatible strains (∼5.2 × 10^−5^ for H53 × RM *ΔhdrB* and ∼3.7 × 10^−5^ for RM *ΔtrpA* × H98). Genome sequencing of 10 hybrids also indicated that they were all *H. mediterranei* which had received genetic material from *H. volcanii*, indicating that genetic transfer during mating always occurred in one direction (from *H. volcanii* to *H. mediterranei*)(Naor *et al*., 2012). It is possible that the presence of RmeRMS in *H. volcanii* may prevent the transfer of genetic material from *H. mediterranei* to *H. volcanii* due to exposed RmeRMS sites. It is also possible that the lower interspecies mating efficiency is due to *H. mediterranei* containing CRISPR spacers which target the genome of *H. volcanii*, since interspecies mating efficiency is further reduced when a CRISPR is added to *H. mediterranei* which is specifically targeted by *H. volcanii* (Turgeman-Grott *et al*., 2019). Future interspecies mating experiments in which *H. volcanii* ΔRM derivative strains are crossed with *H. mediterranei* could elucidate the impact of RM systems on mating efficiencies between these two species.

Although the Halobacteria are highly recombinogenic, distinct phylogroups have been observed even within the same geographic location (Papke *et al*., 2007; Fullmer *et al*., 2014). This suggests that there are barriers to recombination within the Halobacteria which limit interactions between phylogroups and allow for speciation to occur. Our results indicate that when RM systems are incompatible between mating strains of *H. volcanii*, there is a decrease in mating efficiency. This RM-driven decrease in recombination suggests that strains of Halobacteria which have incompatible sets of RM systems might recombine less frequently in natural environments, resulting in the eventual divergence of incompatible strains. However, the results also indicate that RM systems are not a strong barrier to gene flow and are not very efficient in preventing mating between RM-incompatible strains, similar to CRISPR-Cas, glycosylation, and archaeosortases.

## Author contributions

Conceptualization, R.T.P., A.M.M, and M.O.; Funding Acquisition, R.T.P. and J.P.G.; Formal Analysis, M.O., A.M.M. and A.S.L.; Methodology, M.O., A.M.M., and R.T.P; Writing—Original Draft, M.O. and R.T.P.; Writing—Review & Editing, M.O., A.M.M., A.S.L., R.T.P., and J.P.G.

